# Species- and strain-level assessment using *rrn* long-amplicons suggests donor’s influence on gut microbial transference via fecal transplants in metabolic syndrome subjects

**DOI:** 10.1101/2020.09.11.292896

**Authors:** Alfonso Benítez-Páez, Annick V. Hartstra, Max Nieuwdorp, Yolanda Sanz

## Abstract

Fecal microbiota transplantation (FMT) is currently used for treating *Clostridium difficile* infection and explored for other clinical applications in experimental trials. However, the effectiveness of this therapy could vary, and partly depend on the donor’s bacterial species engraftment, whose evaluation is challenging because there are no cost-effective strategies for accurately tracking the microbe transference. In this regard, the precise identification of bacterial species inhabiting the human gut is essential to define their role in human health unambiguously. We used Nanopore-based device to sequence bacterial *rrn* operons (16S-ITS-23S) and to reveal species-level abundance changes in the human gut microbiota of a FMT trial. By assessing the donor and recipient microbiota before and after FMT, we further evaluated whether this molecular approach reveals strain-level genetic variation to demonstrate microbe transfer and engraftment. Strict control over sequencing data quality and major microbiota covariates was critical for accurately estimating the changes in gut microbial species abundance in the recipients after FMT. We detected strain-level variation via single-nucleotide variants (SNVs) at *rrn* regions in a species-specific manner. We showed that it was possible to explore successfully the donor-bacterial strain (e.g., *Parabacteroides merdae)* engraftment in recipients of the FMT by assessing the nucleotide frequencies at rrn-associated SNVs. Our findings indicate that the engraftment of donors’ microbiota is to some extent correlated with the improvement of metabolic health in recipients and that parameters such as the baseline gut microbiota configuration, sex, and age of donors should be considered to ensure the success of FMT in humans.

**Trial registration:** The study was prospectively registered at the Dutch Trial registry - NTR4488 (https://www.trialregister.nl/trial/4488).

## Introduction

Microbiome research advances rapidly and the study of microbial communities demands the implementation of sequencing strategies permitting the identification of microbial species and strains of complex ecosystems and allowing tracing of microbes engraftment in clinical trials evaluating microbiome replacement strategies. Among them, fecal microbiota transplantation (FMT) is widely recognized as an efficient therapy for *Clostridioides difficile* infection that outperforms antibiotic treatment ^1^. Despite the potential risk for the transmission of infectious diseases by FMT, this approach has shown efficacy in ameliorating intestinal inflammation and metabolic disorders, such as insulin resistance ^2-4^. The mode of FMT action is thought to depend on the engraftment of live microbes from donors to recipients, although some clinical trials leave the door open for hypotheses about alternative mechanisms ^5^. The demonstration that the disease amelioration is linked to the ability of donor’s microbes to thrive in the recipient gut is challenging but necessary to ascertain whether particular microbes play a therapeutic role in FMT trials. Metagenomics based on DNA shotgun sequencing is currently the only strategy that enables us to reach such a resolution level, but it is still very expensive, and few strain profiling results adopting such protocols have been provided in FMT studies ^6-8^. Most of FMT microbiome surveys have been performed through the targeted amplification of universal gene markers, mainly including a few hypervariable regions (e.g., V3 and/or V4) of the bacterial 16S rRNA gene ^9, 10^. This technique shows limitations since it only allows changes affecting the global microbiome structure to be captured and does not permit precise taxonomic identifications at the species level, mostly because of the limited resolution of this gene marker in discriminating closely related microbes. This issue is of particular relevance to accurately trace species engraftment in FMT or demonstrate microbe colonization in probiotic-based clinical trials. We previously developed a Nanopore-based amplicon sequencing method to improve the resolution of taxonomic identifications by studying the microbial genetic variability of nearly complete 16S rRNA genes ^11^. This approach has been demonstrated to perform well in a wide variety of microbiota studies ^12-15^. More recently, we have also pioneered a new methodology combining the sequencing of an extremely variable multilocus region with sample multiplexing ^16^ to improve the species-level characterization of complex microbial communities ^12, 17^. The implementations mentioned above of nanopore-based technology and their multiple applications to assess microbial communities allow us to gain knowledge in the configuration of different analytical pipelines improving the processing of such data as updated protocols are delivered ^18-20^.

We have made a great effort to compile a great deal of genetic information on *rrn* sequences from thousands of bacterial species (*rrn_db*, freely available at https://github.com/alfbenpa/rrn_db) ^16^. Nevertheless, the limited number of properly annotated DNA sequences available in public repositories of bacteria from multiple ecosystems restricts the utilization of this approach in less explored habitats ^12, 21^. Notwithstanding, the utility of *rrn*-based analysis for surveying biological diversity has been explored across kingdoms, showing promising results in the identification of metazoans by metabarcoding approaches ^22^.

Despite the issues concerning unlinked 16S rRNA and 23S rRNA markers ^23^ that can be addressed in certain bacterial groups and environmental samples via the study of individual regions, we think the long-read amplicon sequencing of bacterial *rrn* regions could help fill the existing gap to demonstrate the transfer and stable colonization of microbes under FMT strategies in a cost-effective manner.

In this study, we aimed to evaluate the donor’s influence on the transference of specific intestinal microbes to FMT recipients using a unique-donor-to-multiple-recipients approach and amplicon sequencing of the nearly complete *rrn* regions using Nanopore-based sequencing in a trial evaluating the ability of FMT to alleviate metabolic syndrome. We obtained *de novo* information on the *rrn* region from predominant bacterial species inhabiting the human gut, enriching the *rrn* existing database with the annotations of dozens of microbial species and strains. We assessed to what extent the FMT changed the gut microbiota composition of recipients at the species level and, more importantly, tracked the strain-level variation to investigate microbiota engraftment. Overall, our results demonstrated a specific microbial transference between donors and recipients, which likely mediated health effects in metabolic syndrome patients, and that this transference partly depends on the donor-recipient pair, their baseline gut microbiota configuration, and certain donor-associated demographic variables.

## Results

### Assembly of human gut microbiota-derived rrn sequences

The evaluation of the gut microbiota in subjects with metabolic syndrome submitted to FMT aimed at revealing microbiota engraftment; it required the comprehensive analysis of Nanopore sequencing data to generate reliable genetic information supporting species- and strain-level variation additionally to that compiled in our previous *rrn* repository ^16^. During the first step of *rrn* assembly (Figure 1), based on mapping forward reads against the nonredundant NCBI 16S database and the selection of high-quality alignments, we detected the presence of 381 different species. However, preliminary *rrn* assembly was initiated only for those species with at least 10X coverage (250 in total), supporting high-quality consensus *rrn* sequences in downstream analyses. The mapping of reverse reads against the preliminary *rrn* assemblies permitted us to confirm the detection of 229 microbial species. Full *rrn* sequences assembled with reliable taxonomy annotations were retrieved disregarding their individual coverage level (e.g. low, intermediate, or high- Figure S1).

**Figure 1.**
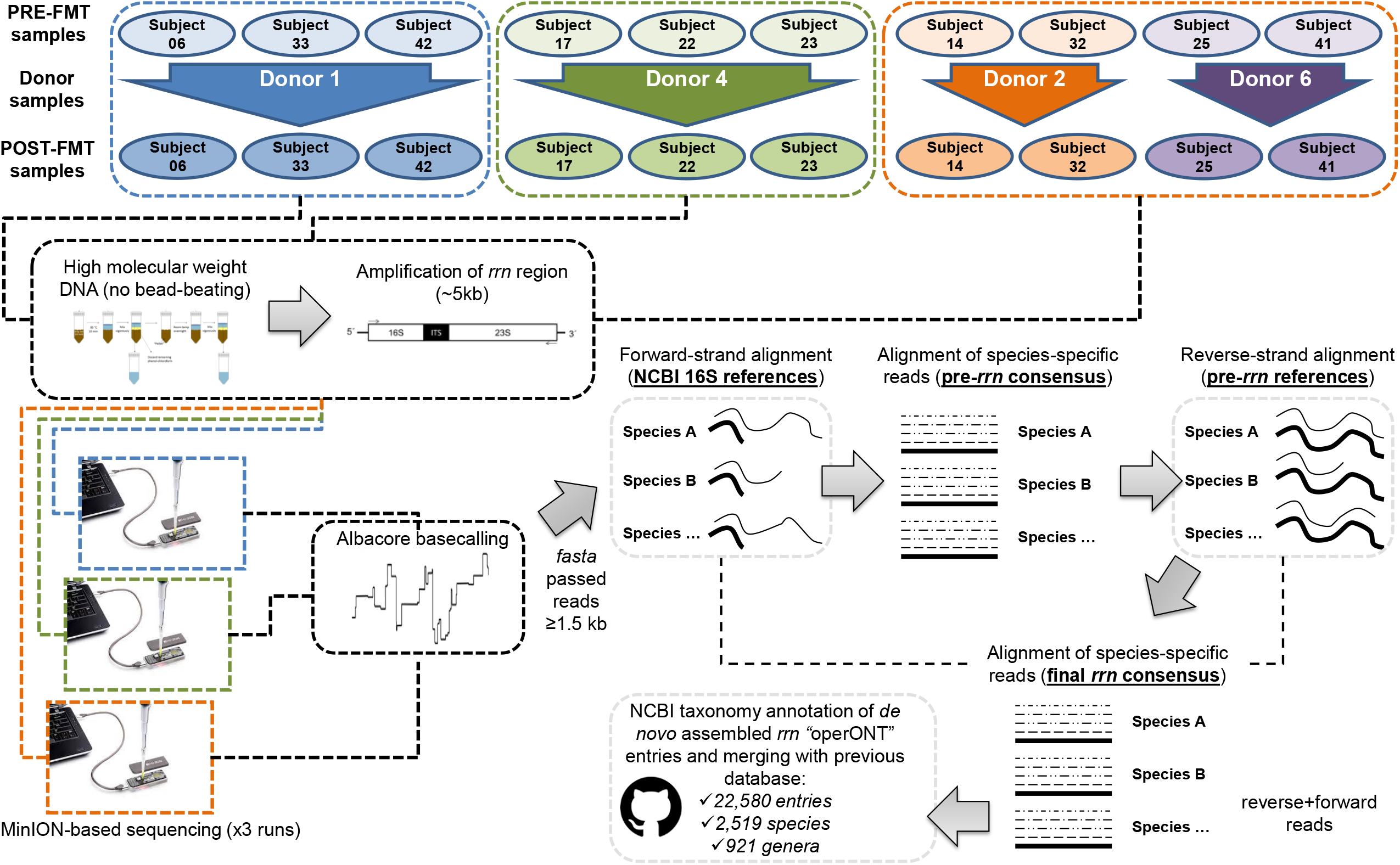
Graphical description of the study. Data acquisition and processing steps, including sample selection, amplicon sequencing, and the general pipeline for the *de novo* assembly of rrn regions from human gut microbiota, are depicted.

### Species abundance shifts associated with FMT

Sample-derived forward and reverse reads were mapped to assess shifts in diversity and taxonomic features. Among all reads, only 43% supported alignments with the highest quality (according to identity and length) and were used in downstream analyses. After taxonomic assignment, we found no large changes in any of the alpha diversity descriptors analyzed. Nevertheless, we found that FMT increased the richness (Chao’s index) of the microbiota in six out of the ten recipients (Figure S3), which showed a gain of 31 species on average. In the remaining four recipients, there was a reduction in richness resulting in an average loss of 16 species as a result of FMT. The beta diversity evaluation based on the Bray-Curtis dissimilarity index indicated no major shifts in the microbial structure of recipients as a consequence of FMT (PERMANOVA = 1.02, *p* = 0.403). Nevertheless, the gut microbiota composition of the recipients was strongly influenced from greater to lesser extents by the sequencing run (PERMANOVA = 3.69, *p* = 0.001), sex (PERMANOVA = 1.70, *p* = 0.032), and donor (PERMANOVA = 1.69, *p* = 0.004). The results of this multivariate analysis are shown in Figure 2A. Globally, we observed that POST-FMT samples tended to be mapped closer to those from donors. To further assess this hypothesis, we compared the distances (Bray-Curtis weighted metrics) between the respective donors and the PRE-FMT microbiota or the POST-FMT microbiota. As a result, we noted that donor microbiota-PRE-FMT pairs had a trend to be more dissimilar than the donor microbiota-POST-FMT pairs, as indicated by a decrease in the Bray-Curtis distance (PRE-FMT vs Donor median = 0.79, POST-FMT vs Donor median = 0.71, average delta = 0.09, *p* = 0.075) (Figure 2B).

**Figure 2.**
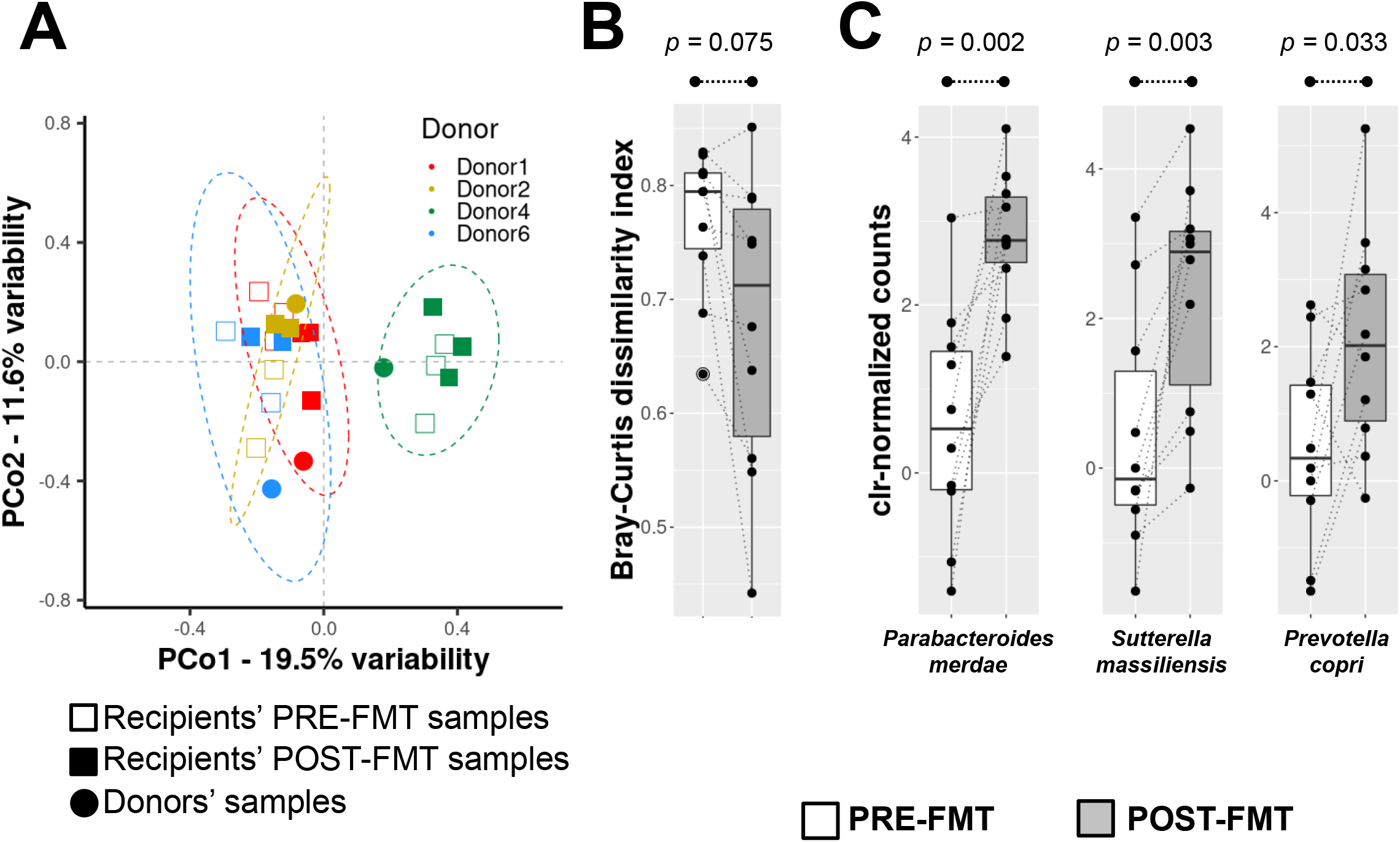
Beta diversity of the microbial communities assessed by *rrn* sequencing. A - Scatter plot compiling data from the multivariate analysis (principal coordinate analysis - PCoA) of the microbiota from recipients and donors participating in FMT. The donor and recipient samples and the sampling time points are defined according to the legend at the top. PCo; principal coordinate (the two most informative coordinates are shown). Color legend corresponds with the distribution of the different PRE-FMT and POST-FMT samples into the groups linked to the four donor samples evaluated (see legend into the plot). B - A Genetic distance-based approach for evaluating microbiota transfer between donor and recipient pairs. The microbial community structures of the PRE-FMT and POST-FMT samples were compared with those of the donors through the calculation of the Bray-Curtis dissimilarity index and are represented as boxplots. The Wilcoxon signed-rank test for paired samples was used to compare differences in genetic distances. C – Microbiota abundance shifts after FMT. Abundance data distribution for three species selected from those described in Table 1 with meaningful increase after FMT is shown in boxplot fashion.

We performed an LMM analysis, which provided similar results to the beta diversity evaluation. We found that sequencing batch, sex, and donor were the main covariates influencing the microbiota data. Furthermore, we found that age and baseline BMI also explained the gut microbiota variation among the subjects of this study to some extent. After including the variables mentioned above as random effects in the model, a list of microbial species showing alterations as a consequence of FMT was retrieved (Table 1), and some of them exhibiting an increased abundance as a consequence of the FMT are plotted in Figure 2C. Ten different microbial species were found to be differentially increased when paired samples obtained before and after FMT were compared, and only three of them (e.g. *Bifidobacterium adolescentis*) seemed to show a decline because of transplantation (Table 1).

**Table 1.** Changes in abundance and prevalence of gut microbial species after FMT.

| Species | Abundance<br>PRE-FMT<br>(N = 10)<sup>1</sup> | Abundance<br>POST-FMT<br>(N = 10)<sup>1</sup> | LMM<br>estimate<sup>2</sup> | Prevalence<br>PRE-FMT<br>(N = 10) | Prevalence<br>POST-FMT<br>(N = 10) | p-value |
| --- | --- | --- | --- | --- | --- | --- |
| *Parabacteroides merdae* | 0.58 ± 1.37 | 2.80 ± 0.79 | 2.07 | 100% | 100% | 0.002 |
| *Prevotella copri* | -0.04 ± 0.50 | 0.90 ± 0.57 | 1.35 | 90% | 100% | 0.033 |
| *Sutterella massiliensis* | 0.44 ± 1.61 | 2.36 ± 1.53 | 1.64 | 90% | 100% | 0.003 |
| *Paraprevotella clara* | -0.31 ± 1.14 | 1.18 ± 1.26 | 1.49 | 70% | 100% | 0.016 |
| *Alistipes shahii* | 1.36 ± 0.87 | 2.27 ± 0.66 | 1.05 | 90% | 100% | 0.005 |
| *Butyricimonas virosa* | -0.38 ± 0.86 | 0.43 ± 0.92 | 1.01 | 80% | 100% | 0.013 |
| *Bacteroides eggerthii* | -0.03 ± 0.89 | 0.50 ± 0.57 | 0.49 | 100% | 100% | 0.029 |
| *Clostridium saccharolyticum* | -0.03 ± 0.84 | -0.54 ± 0.76 | -0.57 | 80% | 100% | 0.076 |
| *Bifidobacterium adolescenti* | 0.38 ± 0.54 | -0.50 ± 0.91 | -0.92 | 70% | 70% | 0.027 |
| *Phascolarctobacterium faecium* | 1.64 ± 1.96 | 0.51 ± 1.45 | -1.47 | 70% | 90% | 0.058 |

1 Data expressed as the mean of clr-normalized counts ± standard deviation (sd). N, sample size.
2 LMM estimate resulting from the global pattern of abundance in POST-FMT samples with respect to baseline information (PRE-FMT samples). Species altered as a consequence of FMT ordered by the size effect (LMM estimate).

### Transferred and engrafted gut microbial species

To assess species engraftment, we next designed a proof of concept analysis to discriminate strain-level variation and thus trace microbes engraftment more accurately. For this purpose, we selected *Parabacteroides merdae*, a predominant species in the samples assessed that showed the greatest increase as a result of FMT (Figure 3A), and *Faecalibacterium prausnitzii*, a highly abundant species in the analyzed samples for which evident transfer between the donor-recipient pairs was not observed (Figure 3B), to detect single-nucleotide variations (SNVs) indicative of strains identity/transference. After a massive analysis of the total dataset merging the species-specific *rrn* reads derived from all subjects (see methods), we identified three potential informative SNV sites in the *P. merdae rrn*, whereas six were found in the *F. prausnitzii* counterpart (Figure 3C-D). In an attempt to explore the natural existence of such genotypic variation inferred from our analysis in a predictive manner, we looked for the presence of similar variants in the NCBI non-redundant nucleotide database using the 100 nt surrounding region as a query for all *P. merdae* SNVs. The inferred genetic variability was replicated at least for SNV-3527, where multiple matches presented A and G nucleotides at this position (Figure S4). When we studied the nucleotide frequencies of these particular SNVs across samples, a clear transfer pattern of the *P. merdae* genotype from donor to recipient was detected, as indicated by the decreased genetic distance (Bray-Curtis index for assessing the abundance of the four nucleotides in all SNVs at once, one-sided Wilcoxon Signed-Rank test = 11, *p* = 0.051) between donor samples and POST-FMT compared to that of POST-FMT and PRE-FMT pairs (Figure 3E). By contrast, the genetic distances retrieved after the comparison of *F. prausnitzii* genotypes between POST-FMT or PRE-FMT samples and their donors did not indicate the transfer of strains belonging to this species from donors to recipients (Figure 3F). Comparative analysis of the delta Bray-Curtis distance (POST vs DONOR minus PRE vs DONOR samples) between both species also suggests that only *P. merdae* was transferred from donors to recipients, since the distance between POST-FMT and donor samples tended to decrease (Figure 3G, *p* = 0.056). The direct Sanger sequencing of *P. merdae* SNV-1254, SNV-2293, and SNV-3527 in the two recipients and their common donor supported the results of our long-read-based assessment of FMT (Figure 3H). Globally, we observed the presence of a mixture of strains in some of the samples analyzed, given the basecalling profile visualized in the electropherograms (e.g., SNV-3527 of Donor1 and 06-Pre samples). This pattern of strain coexistence was more evident in the POST-FMT samples of both analyzed recipients (see SNV-2293 and SNV-3527 in Figure 3H). The predominant haplotype observed in Donor1 (A1254-C2293-A3527) was transferred to the two recipients explored and became dominant after FMT. Similar patterns of transfer were observed in other donor-recipient pairs. Globally, our results demonstrated that the increased abundance of *P. merdae* after the FMT was a direct consequence of donor strain transmission. This is likely the case for other bacterial species that increased as a result of FMT.

**Figure 3.**
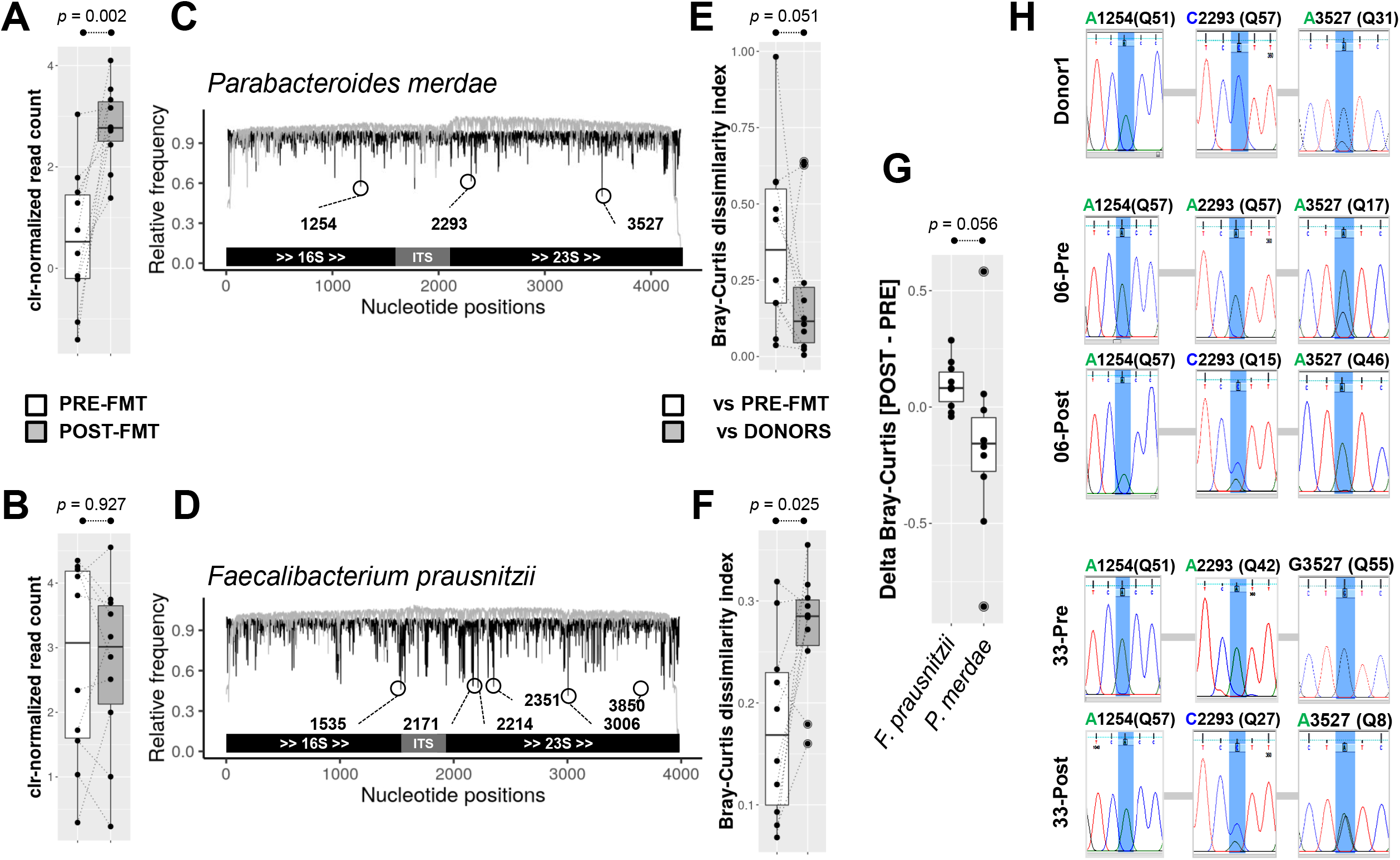
Single-nucleotide variation (SNV) analysis to detect species transfer and engraftment. The species *Parabacteroides merdae* and *Faecalibacterium prausnitzii rrn*s were used to reveal SNVs within each species and their abundance as a consequence of FMT (A and B panels, respectively). Variation in one unit of clr-normalized counts is roughly equal to a doubling in raw DNA read counts according to data distribution in both cases. C and D - The mapping, indexing, and pileup of thousands of reads of *P. merdae* and *F. prausnitzii rrns* allowed us to identify polymorphic sites exhibiting an even frequency for at least two nucleotides. These potential SNVs are highlighted inside open circles, and the position number in the *rrn* sequence is also indicated. The structural arrangement of *rrn* is illustrated on the x-axis. The black lines indicate the relative frequency of the dominant allele, whereas the gray lines indicate the relative coverage per site related to the average across the *rrn*. E and F - The distribution of the Bray-Curtis dissimilarity index of the microbiota between POST-FMT and PRE-FMT samples and the corresponding donors, using the combined nucleotide frequencies of SNVs detected in *P. merdae* and *F. prausnitzii*, respectively. The Wilcoxon signed-rank test for paired samples was used to compare the genetic distances of the microbiotas. G – Assessment of delta Bray-Curtis distances computed as POST-FMT vs DONOR samples minus PRE-FMT vs DONOR samples for *Faecalibacterium prasunitzii* and *Parabacteroides merdae* species. H - Electropherograms obtained from Sanger sequencing for the strains of *P. merdae* SNV-3527, SNV-2293, and SNV-1254. The predominant alleles inferred for every sample (based on the Q-score of base calling) supported the hypothesis that the strains were transferred between donor and recipient pairs, as anticipated during the Nanopore-based assessment.

### Correlation between clinical variables and species abundance after FMT

Among the multiple clinical variables evaluated in these subjects with metabolic syndrome, FMT tended to induced positive changes in the markers of glucose metabolism and blood pressure in the recipients (Table 2). Fasting insulin (*p* = 0.030), fasting glucose (*p* = 0.074), and consequently HOMA-IR (*p* = 0.005) tended to improved in the recipients after FMT. In line with these findings, there was an apparent decrease in the glycosylated hemoglobin concentration after FMT (*p* = 0.060). On the other hand, systolic and diastolic blood pressures were lowered as a consequence of donor FMT treatment, being reduced by 10% (*p =* 0.028) and 19% (*p* = 0.0008), respectively (Table 2). The concentrations of fecal SCFAs, such as fecal butyrate (*p* = 0.018) and acetate (*p* = 0.033), were decreased after FMT, whereas that of propionate was not (*p* = 0.280). Regarding markers of lipid metabolism, HDL cholesterol levels in plasma were reduced (*p* = 0.032). By computing rank-based Kendall’s *τ* (tau) parameter, we established associations between these clinical variables and the alterations in the abundance of microbial species as a consequence of FMT (see Table 1). Species such as *P. merdae*, which was increased after FMT, were negatively correlated with diastolic blood pressure (τ = -0.44 and FDR = 0.062), whereas *A. shahii* was negatively correlated with the abundance of fecal acetate and butyrate (τ = -0.50, -0.42 and FDR = 0.017, 0.042, respectively). The above correlations were not detected for other closely related and abundant species (e.g., *Parabacteroides johnsonii, Parabacteroides distasonis, Parabacteroides goldsteinii, Alistipes finegoldii, Alistipes onderdonkii, Alistipes putredinis* or *Alisti*pes *indistinctus*).

**Table 2.** Clinical variables altered after FMT.

| Clinical outcome | PRE-FMT<br>(N = 10)<sup>1</sup> | POST-FMT<br>(N = 10)<sup>1</sup> | Statistics |
| --- | --- | --- | --- |
| women = 5 (50%) |  |  |  |
| age (years) = 60.2 ± 6.4 |  |  |  |
| BMI (kg/m<sup>2</sup>) = 32.9 ± 3.5 |  |  |  |
| **Diastolic blood pressure** | 87.4 ± 8.2 | 72.0 ± 9.9 | $v = -16.9, p < 0.001$ |
| **Systolic blood pressure** | 138.6 ± 15.3 | 124.5 ± 14.2 | $v = -14.2, p = 0.028$ |
| **Fasting glucose (mmol/L)** | 5.49 ± 0.35 | 5.31 ± 0.50 | $v = -0.18, p = 0.074$ |
| **Fasting insulin (mg/dL)** | 84.7 ± 30.9 | 63.6 ± 21.9 | $v = -19.0, p = 0.030$ |
| **HOMA-IR** | 2.96 ± 0.99 | 2.16 ± 0.78 | $v = -0.93, p = 0.005$ |
| **Hba1c (mmol/L)** | 36.5 ± 4.1 | 35.7 ± 3.6 | $v = -0.96, p = 0.060$ |
| **Plasma HDL (mmol/L)** | 1.59 ± 0.29 | 1.40 ± 0.23 | $v = -0.191, p = 0.032$ |
| **Fecal butyrate (μmol/g)** | 88.6 ± 44.2 | 60.3 ± 40.9 | $v = -31.6, p = 0.018$ |
| **Fecal acetate (μmol/g)** | 434.0 ± 148.3 | 308.2 ± 121.1 | $v = -135.7, p = 0.033$ |

1 Data expressed as the mean ± standard deviation (sd). N, sample size.
$v$ = variation between groups analyzed by applying an LMM (PRE-FMT group as reference), HDL = high-density lipoprotein, Hba1c = glycosylated hemoglobin.

### Donors’ influence on species engraftment and amelioration of metabolic dysfunction

After showing that microbiota engraftment took place between donor and recipients, and that reversion of certain metabolic alterations may have been guided by particular microbial species transferred from donors to recipients, we wanted to explore to what extent the donors and their microbiota configurations can explain the FMT success. For such an aim, we ranked the metabolic improvement individually (from 1 to 10, from the lowest to the highest), including the reduction of blood pressure, fasting glucose and insulin levels, glycosylated hemoglobin, HOMA-IR, weight loss, and BMI of the ten subjects of the FMT. Then, the health score resulting from the sum of all scores for every subject was estimated. This score was related to t the dissimilarity index (Bray-Curtis) between the baseline microbiota configuration of donor-recipient pairs and the community structural shift resulting from the FMT in every recipient, taking into account their donor pair (Figure 4). Globally, we observed that metabolic health scores correlate to some extent with the baseline dissimilarity between donors and recipients microbiotas and the community structural changes after FMT. Consequently, the more dissimilarity in the baseline gut microbiota of donors and recipients, the lower the benefits to revert the metabolic dysfunction in recipients and less impact on their gut microbiota structure after FMT. The above observation applies to almost all donor-recipient pairs, except for those including the Donor2. Although we observed a linear relationship between the averaged dissimilarity between donor-recipient pairs and the respective shift in the community structure after FMT (adjusted R-squared: 0.956, p = 0.013), a rank-based approach (Kendall’s tau) showed those parameters were not well correlated with the metabolic health score (τ ≥ -0.24, p ≥ 0.381). Therefore, additional traits yet to explore in donor-recipient pairs should help in the selection of subjects for succeeding in FMT. In line with this hypothesis, we finally evaluated the relationship between the metabolic health score and the recipients sex, showing that women exhibited a larger improvement in metabolic parameters than men (women [N = 5] = 49.6 ± 13.8 health score, men [N = 5] = 33.6 ± 11.3 health score, *t*-test = 6.62, p = 0.003). Moreover, we found a strong rank-based correlation between the age difference of donor-recipient pairs (donor age - recipient age) and the metabolic health score (Kendall’s τ = -0.51, p = 0.046), suggesting that the youngest the donor (with respect to the recipient), the greater reversion of metabolic dysfunction.

**Figure 4.**
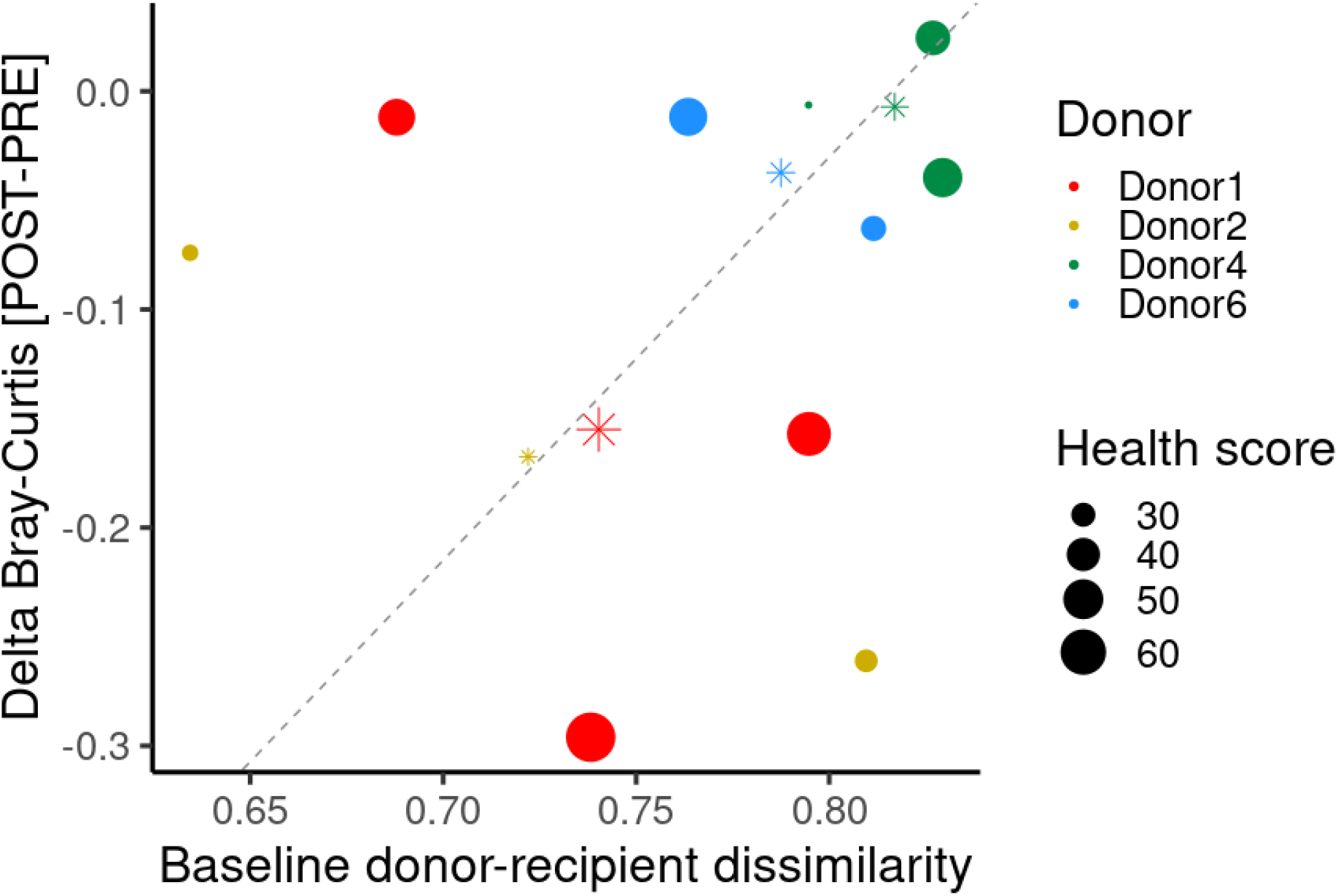
Donors’ influence on FMT success. A scatter plot showing the comparison of gut microbiota between donor-recipient pairs at baseline (x-axis, Bray-Curtis dissimilarity index computed) versus the shift in the microbiota community structure of every recipient before and after the FMT (y-axis). The size of filled dots is determined by the sum of ranks linked to the improvement of all metabolic parameters evaluated (blood pressure, fasting glucose and insulin, glycosylated hemoglobin, HOMA-IR, weight loss, and BMI loss) (metabolic health score). Color legend is associated with donor information. Asterisks and size represent the average values of all recipients linked to every donor. Dashed grey line indicates linear correlation based on averaged values (asterisks).

## Discussion

Here, we explored the *rrn* amplicon sequencing of the human gut microbiota through Nanopore sequencing and the *de novo* assembly of sequences of this multilocus hypervariable region to gain insights into the corresponding genetic markers and to demonstrate their potential use for deeply characterizing complex communities at the species and strain levels, permitting the assessment of microbe engraftment between donors and recipients in an FMT clinical trial. The utility of *rrn* operon sequencing via the MinION™ platform has already demonstrated a notable performance in resolving microbial communities at the species level, with the potential to define strain variability for particular species ^24, 25^.

We provided *de novo* reliable assemblies for more than two hundred *rrn* region sequences of human gut microbes. We were able to reconstruct a large proportion of *rrn* regions from approximately 40 new microbial species, and more than 150 new strains were absent in the first database release ^16^. These numbers, limited mainly by sequencing coverage, could also be reduced by the DNA extraction method used, in which a physical disruption step was omitted to maintain DNA integrity and ensure the successful amplification of the 5 kbp amplicons. However, the homogeneous processing of all samples resulted in comparable microbial profiles before and after FMT and did not affect the species-specific evaluation conducted in this study.

By using a dataset consisting of approximately 400 K Nanopore reads derived from 24 samples, employing the updated *rrn* database, and controlling the covariates influencing the microbiota, we revealed changes in the human gut microbiota at the species and strain levels as a consequence of an FMT intervention. This long amplicon-based approach enabled us to detect increases in species such as *Parabacteroides merdae, Butyricimonas virosa, Prevotella copri*, and *Sutterella massiliensis* as a result of FMT. In addition to age, sex, sequencing runs (recognized stochastic run-to-run variation confounding the interpretation of microbiota data ^26, 27^), and baseline BMI, we found that the donor was a critical covariate contributing to the impact of FMT on the recipient microbiota when using a unique donor for multiple recipients. These results are in agreement with those previously published on the basis of short reads from the V4 hypervariable region of the 16S rRNA bacterial gene explored in a larger cohort of subjects ^3^, for instance, the significant increase of *P. copri* after FMT; however, our results outperform the taxonomic resolution achieved previously, thus providing a more accurate gut microbiota survey. Interestingly, a recent FMT study on the treatment of ulcerative colitis (UC) in which the microbiota was analyzed by sequencing short reads also showed increases in *Sutterella* species as a consequence of the intervention ^28^. Similarly, the abundance of *Butyricimonas* species was shown to be increased as a consequence of an FMT intervention aimed at eradicating antibiotic-resistant bacteria ^29^. These findings suggest that some species may frequently show changes as a result of FMT in humans, regardless of the condition of the recipient. Nevertheless, the presumable transfer of species between donors and recipients and their engraftment could not be confirmed in those studies based on short-read amplicon technology due to the lack of sufficient resolution. Additionally, potential replacement with closely related species in the recipient could be impossibly monitored in conventional microbiota surveys based on short reads. To shed light on the ability of the *rrn* sequencing approach to assess SNVs that are likely associated with strain diversity, thus demonstrating donors’ microbe engraftment in recipients, we designed a proof of concept analysis selecting *P. merdae*, a species that exhibited a remarkable increase in the recipients’ gut after FMT, and *F. prausnitzii*, which seemed to not be affected by FMT intervention. The combined information on different informative SNVs from the *P. merdae rrn* sequence clearly suggests that the increase in this species observed in recipients after undergoing FMT was likely because of strain transfer from the donors. This inference is further supported by the existence of nucleotide variants inferred from Nanopore data processing in either the samples evaluated in the present study or the genomic data present in public repositories and derived from different sequencing methodologies.

The taxonomic resolution of our sequencing approach, proving the transference of specific bacterial strains from donors to recipients, helped draw partial community structures at the species level and determine compositional distances and their shifts as a consequence of the FMT. This strategy helped to more firmly support a causal relationship between the changes in the gut microbiota and the improvements in the recipient’s clinical variables. In this regard, we attempted to demonstrate that overall improvement of metabolic health (global score) was associated with baseline dissimilarity between microbial communities of donors and recipients and with their microbiota configuration shifts after FMT. We observed that the more similar is the gut microbiota of the donor-recipient pairs at baseline, the better is the microbiota transference and the improvement in metabolic parameters. However, this correspondence was not proven for all pairs; possibly, additional traits of donors and recipients yet to be determined need to be considered for an optimal selection of donors to ensure the clinical success of the FMT. One of those unknown additional factors could be the sex of the recipient and donor. Indeed, we observed that women recipients exhibited higher metabolic health score after FMT than men. Given that the sex of donors and recipients was matched, further studies with a mixed-sex distribution would be needed to determine i) if women’s fecal material can be better engrafted regardless of the recipient sex, or ii) whether women respond better to FMT irrespective of the sex origin of the FMT material. We have previously unveiled differential responses of women and men gut microbiota to a particular dietary intervention, suggesting that such disparate sex-associated microbial configurations and related traits could plausibly explain the success of FMT ^30^. Nonetheless, previous FMT interventions in the context of IBS show no differential response between women and men ^31^. Finally, an individual-specific response to FMTs cannot be disregarded as in our trial the material from Donor1 exerted the most positive impact on the metabolic health of their three different recipients. Consequently, the role of the personal features of the donor, particularly those influencing the gut microbiota composition and metabolic potential (e.g. diet, lifestyle) as well as other host covariates (e.g. age) linked to successful FMTs should be further investigated to better guide donor selection and identify microbial signatures improving metabolic health.

## Conclusions

The molecular methods and data analyses presented here enabled reliable microbial surveys at the species level and the inference of strain variations, taking into account the most abundant human intestinal bacterial species. To the best of our knowledge, this is the first microbiota assessment of fecal samples derived from donors and recipients involved in a FMT clinical trial to demonstrate transference and engraftment of specific bacterial species by exploring the genetic diversity at the strain level using amplicon sequencing. The genetic and taxonomy information retrieved from advanced sequencing methods used here permitted us to evaluate the donors’ influence on the gut microbiota transference and clinical success of the FMT. The determinants of the success include a high relatedness between the donor-recipient baseline microbiota configurations, the sex and the age of the donor, which seem to influence the engraftment of intestinal microbes and the amelioration of metabolic syndrome markers. Further studies should be warranted to confirm our findings in larger intervention trials and demonstrate their predictive value.

## Materials and Methods

### Subjects, samples, and clinical data

Samples were obtained upon receiving informed consent from participants in a previous FMT clinical trial carried out under the framework of the MyNewGut project (http://www.mynewgut.eu/) between September 2014 and March 2017. The details of the study design are publicly available elsewhere ^3^. Briefly, a total of twenty-four fecal samples were analyzed in the present study. Twenty samples were obtained from 10 recipients undergoing allogenic FMT, who provided one sample before (PRE-FMT fecal samples) and one sample 4 weeks after (POST-FMT fecal samples) the intervention. The samples of 4 donors were also analyzed, and the effect of this variable on the recipients was considered in the data analysis. This subset of samples was mainly assessed to explore differential species engraftment in multiple FMT recipients with common donors. The sample size evaluated in this study is still representative according to parameters described previously for this intervention arm (two-sided significance level of 0.05 and power of 80%, minimum sample size = 10) ^3^. Normality test was completed prior to execute parametric or non-parametric tests to assess changes in the clinical variables explored.

### DNA extraction, multilocus amplification, and sequencing

Microbial DNA was recovered from 100 mg feces samples by using the QIAamp® Fast DNA Stol Mini kit (Qiagen, Hilden, Germany) according to the manufacturer’s instructions and omitting cell disruption by mechanical methods (bead-beating) to preserve DNA with the high molecular weight needed to amplify amplicons of 5 kbp in length. The *rrn* region comprising the nearly full-length bacterial RNA ribosomal operon (16S, ITS, and 23S) was amplified as previously reported ^16^. Dual-barcoded purified PCR products were mixed in equimolar proportions before sequencing library preparation. In total, three different libraries were prepared from ∼1 µg of mixed amplicon DNA (containing 7, 8, or 10 barcoded samples) using the SQK-LSK108 sequencing kit (Oxford Nanopore Technologies, Oxford, UK) following the manufacturer’s instructions to produce 1D reads. Each library was individually loaded into FLO-MIN107 (R9.5) flow cells (Oxford Nanopore Technologies, Oxford, UK), and sequencing was carried out in a portable MinION™ MkIb sequencer (Oxford Nanopore Technologies, Oxford, UK), operated with *MINKNOW* v1.10.23 software (Oxford Nanopore Technologies, Oxford, UK). Flow cells were primed according to the manufacturer’s instructions, and an ∼18 h run of 1D sequencing (outperforming the 2D approach for this multiplex and amplicon setup, and improving de-multiplexing by using the barcode+primer tags) was then executed for each library and flow cell.

### Data preprocessing

Fast5 files were processed with the *Albacore* v2.1.3 basecaller and default configuration (minimum q-score filter = 7), and *fasta* files were retrieved for downstream analyses. The barcode and primer (forward or reverse) sequence information was used for demultiplexing into the DNA reads generated from forward and reverse primers according to previous procedures ^16^. A size filtering step was configured to retain those reads of least 1,500 nt in length. Barcode and primer sequences were then removed by trimming 50 nucleotides at the 5’ end of forward and reverse reads. The sequencing chemistry and Nanopore data-processing algorithms were the latest available by January 2018, when data was generated. A high-performance basecaller released after the end of this analysis was tested on fast5 data produced getting no significant difference in sequences quality (Figure S4).

### Assembly of rrn regions

The study design and the assembly of *rrn* sequences are graphically explained in Figure 1. This pipeline carries out what we refer to as the strand-wise and reference-assisted method for *rrn* region reconstruction via the following steps:

- Demultiplexing of Nanopore data using barcode and primer sequence information generated two datasets per sample comprising DNA reads led from the 5’
s end of *rrn* amplicons (initiated from forward primer mapping at the 16S rRNA gene start site = forward reads), and DNA reads beginning at the 3’ end of *rrn* amplicons (initiated from reverse primer mapping at the 23S rRNA gene end site = reverse reads).
- Reads from all 24 samples were merged into forward and reverse read subsets.
- Forward read binning into precise microbial species via competitive alignment against the nonredundant 16S NCBI database (release January 2018). The *LAST* aligner ^32^ with a -s 2 -q 1 -b 1 -Q 0 -a 1 -r 1 configuration was used for this purpose. The alignment score was the main criterion for selecting top hits. In the case of multiple hits with the same top score, the alignment was discarded for downstream processing. High-quality alignments were retained according to sequence identity (sequence identity above the 33rd percentile ≥ 85%) and length information (alignments ≥ 1500 nt).
- A maximum of 500 forward reads per species (randomly shuffled), producing high-quality alignments, were selected, and aligned through iterative refinement methods implemented in *MAFFT* v7.310 with the default parameters ^33^, and the consensus sequence was then obtained by using the *hmmbuild* and *hmmemit* algorithms implemented in *HMMER3* ^34^. No significant improvements in consensus *rrn* sequences were obtained using more than 500 sequences per species. Assemblers supporting nanopore data, temporary available at the date of analysis, were tested to produce *de novo* consensus sequences but they failed to produce outputs supporting the expected species-level diversity because of the coverage bias across data.
- Consensus sequences were annotated according to the NCBI taxonomy and used as a reference to bin reverse reads in a similar manner as that used for forward reads. The thresholds for the selection of high-quality alignments based on reverse reads and the preliminary assembly of *rrn* regions were 85% sequence identity (upper 45th percentile) and ≥ 3500 nt in length.
- Reverse reads were compared against the consensus sequences obtained from the forward read assemblies and binned into the respective species according to NCBI taxonomic annotation. A maximum of 500 reverse reads per species (randomly shuffled), producing high-quality alignments, were selected, merged with a maximum of 500 forward reads assigned reciprocally to the same species (randomly shuffled), and aligned through iterative refinement methods implemented in *MAFFT*, after which a new consensus sequence was obtained by using the *hmmbuild* and *hmmemit* algorithms implemented in *HMMER3*.
- Final consensus sequences were annotated according to the concordant NCBI taxonomy, obtained from forward and reverse reads, and used as a reference to study the abundance and prevalence of gut microbiota members at the species level.
- A BLAST-based search of the *rrn* assemblies against the nonredundant nucleotide and reference 16S NCBI databases was performed to evaluate the representation of the reconstructed and identified operons in public databases. The identification of *rrn* assemblies was compared against the SILVA database ^35^ with SINA aligner ^36^. Top hit selection was based on the taxonomy score, TS = Log_10_[alignment score*sequence identity*alignment length].

### Long-read mapping and variant calling

The *rrn* database ^16^ was updated with more than two hundred new *rrn* operon sequences assembled in this study (labeled operONT) and re-annotated to different taxonomic levels according to the NCBI taxonomy database. The mapping of sample reads was performed against *rrn_DBv2* (https://github.com/alfbenpa/rrn_DBv2) by competitive alignment using the *LAST* aligner, as stated above. Taxonomic assignment was based on the best hit retrieved by the calculation of TS (see above), and reliable taxonomic assignments were filtered to those based on alignments with at least 85% sequence identity that were longer than 1500 nt. Species read counts were normalized using the *compositions::clr* R v3.6 function, and a covariate-controlling linear mixed model-based method was used to explore the differential abundance of taxonomic features between conditions as stated in the paragraphs below. Species engraftment was evaluated by the selection of all sample reads mapped to the *Parabacteroides merdae* and *Faecalibacterium prausnitzii* species. To detect informative sites, based on single-nucleotide variants that were probably linked to strain variation, we first proceeded to map all sample reads matching these species to the respective reference *rrn* sequences using *LAST* aligner and the parameters set across this study (see *rrn* assembly). The *maf-convert* algorithm (*LAST* aligner toolkit) was used to generate the respective *SAM* files. The algorithms compiled within *SAMtools* v1.3.1 were used to index, order, and pileup reads as well as to retrieve information regarding the nucleotide frequency per site and coverage (*vcf* files). Site selection was based on *rrn* positions showing no nucleotide dominance (<70% frequency for a single nucleotide), without reductions in the coverage (>75% of the relative coverage), thus obtaining sites with a balanced representation of at least the two predominant alleles likely representing strain variation across all samples. Second, the sample reads that were mapped to the reference species were individually assessed to detect meaningful changes in intestinal bacterial genotypes between the recipients PRE-FMT and POST-FMT samples. The changes in nucleotide frequencies at all selected SNV sites (three for *P. merdae* and six for *F. prausnitzii*) were assessed by calculating the Bray-Curtis dissimilarity index between pairs of POST-FMT and PRE-FMT samples or POST-FMT and corresponding donor samples following statistical evaluation by using the Wilcoxon signed rank test for paired samples.

### Sanger sequencing and validation of Parabacteroides merdae SNVs

The reference *rrn* sequence from *P. merdae* was submitted to the Primer-BLAST web server (https://www.ncbi.nlm.nih.gov/tools/primer-blast/) to retrieve species-specific primer pairs to selectively amplify this *rrn* sequence and primers flanking SNV-1254, SNV-2293, and SNV-3527 for Sanger sequencing. Comparisons against the nonredundant NCBI database and *Parabacteroides merdae* [taxid:46503] as a reference organism were fixed as checking parameters for primer prediction. The *rrn* region from *P. merdae* was amplified via 28 PCR cycles including the following stages: 95°C for 20 s, 61°C for 30 s, and 72°C for 150 s. Phusion High-Fidelity Taq Polymerase (Thermo Scientific) and the pm623 (TGCCGTTGAAACTGGGTTACTTGA) and pm3785 (TTTCGCACAGCCATGTGTTTTGTT) primer pairs were used in the amplification reaction. The PCR products were cleaned with an Illustra GFX PCR DNA and gel band purification kit (GE Healthcare, Chicago, IL, USA) and sequenced by using Sanger technology in an ABI 3730XL sequencer (STAB-VIDA. Caparica, Portugal) using the primers described above (pm623 and pm3785) in addition to the pm1855 (ACCCCTTACGGAGTTTATCGTGGA) and pm2686 (TTCGCGTCTACTCACTCCGACTAT) primers. The sequencing electropherograms from the *ab1* files were visualized with FinchTV v1.4.0 (Geospiza Inc.).

### Diversity and taxonomic analyses at the species level

The prevalence and abundance of a total of 2,519 species contained in the database were evaluated, and diversity analyses were completed taking into account the species with a > 0.01% relative abundance on average (∼250 species in total). Alpha diversity descriptors such as Chao’s index, Shannon’s entropy, Simpson’s reciprocal index, and dominance were obtained by using *QIIME* v1.9.1 ^37^. Similarly, *QIIME* was used to calculate the Bray-Curtis dissimilarity indexes among samples and to perform multivariate exploratory (principal coordinate analysis - PCoA) and statistical (PERMANOVA) analyses. A linear mixed model (LMM – *nlme::lme* R v3.6 function) analysis was also conducted on clr-transformed data to detect differential features in the microbiota before and after the intervention. The inherent variation due to individual features was set as a random effect for each variable analyzed (fixed effect). The possible covariates of the clinical and fecal microbiota that showed differences between the study groups (p-value ≤ 0.05) were selected. Recognized covariates of microbiota, such as age, sex, baseline BMI, and sequencing batch, were identified as significant in this study and included as random effects in the LMM. To identify microbial species potentially linked to clinical variables that were altered as a consequence of FMT, Kendall’s *τ* (tau) parameter was estimated between variable pairs and corrected for multiple testing using the false discovery rate (FDR) approach. Associations were selected according to an FDR p-value ≤ 0.1. Graphics were generated in R v3.6 using the *ggplot2* package.

## List of abbreviations

BMI: body mass index
FDR: false discovery rate
FMT: fecal microbiota transplantation
LMM: linear mixed model
NCBI: National Center for Biotechnology Information
MAG: metagenome-assembled genome
PCoA: principal coordinate analysis
PCR: polymerase chain reaction
RDP: Ribosomal Database Project
rrn: bacterial ribosome RNA operon (16S-ITS-23S)
SNP: single-nucleotide polymorphism
SNV: single-nucleotide variation
UC: ulcerative colitis

## Declarations

### Ethics approval and consent to participate

The study was prospectively registered at the Dutch Trial registry (https://www.trialregister.nl/trial/4488) and was conducted according to the guidelines laid down in the Declaration of Helsinki and the ethical standards of the responsible local committee on human experimentation of the Amsterdam UMC (location AMC) ^3^. The study was registered on August 1^st^, 2014. The first participant was enrolled on September 1^st^, 2014.

### Availability of data and material

The Albacore-basecalled *fast5* files obtained from specific runs are publicly available in the European Nucleotide Archive under accession number PRJEB33947. The updated *rrn* database (*rrn_DBv2*) is publicly accessible at the GitHub repository https://github.com/alfbenpa/rrn_DBv2.

### Authors’ contributions

ABP conceived and designed the study. ABP performed the experimental sequencing research and data analysis, and AVH and MN performed the clinical research. ABP and YS directed the study. ABP and YS wrote the manuscript. All authors reviewed and approved the final version of the manuscript.

## Acknowledgements

The authors thank the Bioinformatics and Biostatistics Unit from Principe Felipe Research Center (CIPF) for providing access to the cluster, co-funded by European Regional Development Funds (ERDF/FEDER) in Valencian Community 2014-2020.

## Figure legends

**Figure S1**. BLAST-based results showing the correct identification of assembled *rrn*s. The top hits supporting the identification are shown for six different *rrn*s assembled with low, intermediate or high coverage (number of reads).

**Figure S2**. Alpha diversity of the recipients’ microbiota before and after FMT. The distributions of the values obtained for four alpha diversity indicators, including Chao’s index, Shannon’s index, the reciprocal Simpson’s index, and the dominance index, are shown as boxplots. The results of the Wilcoxon signed-rank test applied to establish differences between the two groups of samples (POST-FMT and PRE-FMT) are also shown.

**Figure S3**. Searching for SNV genotypic variation in the NCBI nonredundant nucleotide database. The nucleotide sequences spanning positions 1180 to 1280, 2250 to 2350, and 3470 to 3570 of the *P. merdae rrn* operon were used as queries in searches against the nonredundant nucleotide database on the BlastN web server (https://blast.ncbi.nlm.nih.gov/Blast.cgi). The results showed the intrinsic variation attributable to SNV-3527 (as an example) present in genomic records from the NCBI nonredundant nucleotide database including strains with genotype A3527 and others with G3527, both of which were variants inferred from our Nanopore-based sequencing. Nucleotides highlighted in yellow show the 5-mer surrounding the region of SNV-3527, revealed in detail by Sanger sequencing, as shown in Figure 4H. The C2293 genotype was retrieved from a unique matching sequence (CL06T03C08 strain), and A1254 was predominantly found in most of the *P. merdae* strains evaluated (data not shown).

**Figure S4**. FastQC-based assessment of sequences produced by nanopore device. Results from the main basecaller, *Albacore v2*.*1*.*3* and the more recently released high-performance basecaller of nanopore data (*Guppy v5*.*0*.*11*) are shown for all three different sequencing runs generated in this study.

