## Supplementary material for "Species- and strain-level assessment using *rrn* long-amplicons suggests donor’s influence on gut microbial transference via fecal transplants in metabolic syndrome subjects": Figure S1

### ➢*Streptococcus sobrinus* - rrn; length = 4,272 nt; coverage = 10 reads (low)

| Description | Max Score | Total Score | Query Cover | E value | Per. Ident | Accession |
|---|---|---|---|---|---|---|
| Streptococcus sobrinus strain 10919 chromosome, complete genome | 7284 | 43705 | 98% | 0.0 | 97.92% | CP029491.1 |
| Streptococcus sobrinus strain NCTC10921 genome assembly, chromosome: 1 | 7267 | 43629 | 98% | 0.0 | 97.85% | LS483381.1 |
| Streptococcus sobrinus strain NCTC12279 genome assembly, chromosome: 1 | 7267 | 43602 | 98% | 0.0 | 97.85% | LS483378.1 |
| Streptococcus sobrinus strain NIDR 6715-7 chromosome, complete genome | 7267 | 43621 | 98% | 0.0 | 97.85% | CP029560.1 |
| Streptococcus sobrinus strain NIDR 6715-15 chromosome, complete genome | 7267 | 43626 | 98% | 0.0 | 97.85% | CP029559.1 |
| Streptococcus sobrinus strain SL1 chromosome, complete genome | 7267 | 43602 | 98% | 0.0 | 97.85% | CP029490.1 |

### ➢*Acidaminococcus intestini* - rrn; length = 4,120 nt; coverage = 12 reads (low)

| Description | Max Score | Total Score | Query Cover | E value | Per. Ident | Accession |
|---|---|---|---|---|---|---|
| Acidaminococcus intestini RyC-MR95, complete genome | 6968 | 20900 | 98% | 0.0 | 97.56% | CP003058.1 |
| Acidaminococcus fermentans DSM 20731, complete genome | 5729 | 34604 | 98% | 0.0 | 92.16% | CP001859.1 |

### ➢*Eubacterium cylindroides* - rrn; length = 4,081 nt; coverage = 170 reads (mid)

| Description | Max Score | Total Score | Query Cover | E value | Per. Ident | Accession |
|---|---|---|---|---|---|---|
| Eubacterium cylindroides T2-87 draft genome | 7249 | 7249 | 98% | 0.0 | 99.21% | FP929041.1 |
| Faecalitalea cylindroides strain JCM 10261 16S ribosomal RNA, partial sequence | 2697 | 2697 | 36% | 0.0 | 99.14% | NR_113163.1 |

### ➢*Streptococcus thermophilus* - rrn; length = 4,112 nt; coverage = 766 reads (high)

| Description | Max Score | Total Score | Query Cover | E value | Per. Ident | Accession |
|---|---|---|---|---|---|---|
| Streptococcus thermophilus strain GABA chromosome, complete genome | 7476 | 44830 | 98% | 0.0 | 99.83% | CP025399.1 |
| Streptococcus thermophilus strain DGCC 7710 chromosome, complete genome | 7476 | 37381 | 98% | 0.0 | 99.83% | CP025216.1 |
| Streptococcus thermophilus strain ST3, complete genome | 7476 | 44830 | 98% | 0.0 | 99.83% | CP017064.1 |
| Streptococcus thermophilus strain APC151, complete genome | 7476 | 44834 | 98% | 0.0 | 99.83% | CP019935.1 |
| Streptococcus thermophilus strain ND07, complete genome | 7476 | 37381 | 98% | 0.0 | 99.83% | CP016394.1 |
| Streptococcus thermophilus strain KLDS SM, complete genome | 7476 | 44847 | 98% | 0.0 | 99.83% | CP016026.1 |
| Streptococcus thermophilus strain MN-BM-A02, complete genome | 7476 | 37200 | 98% | 0.0 | 99.83% | CP010999.1 |
| Streptococcus thermophilus strain SMQ-301, complete genome | 7476 | 44819 | 98% | 0.0 | 99.83% | CP011217.1 |
| Streptococcus thermophilus ASCC 1275, complete genome | 7476 | 37381 | 98% | 0.0 | 99.83% | CP006819.1 |
| Streptococcus thermophilus JIM 8232 complete genome | 7476 | 44810 | 98% | 0.0 | 99.83% | FR875178.1 |
| Streptococcus thermophilus LMD-9, complete genome | 7476 | 44819 | 98% | 0.0 | 99.83% | CP000419.1 |
| Streptococcus thermophilus strain EPS chromosome, complete genome | 7472 | 44830 | 98% | 0.0 | 99.80% | CP025400.1 |

### ➢*Akkermansia muciniphila* - rrn; length = 4,306 nt; coverage = 1000 reads (high)

| Description | Max Score | Total Score | Query Cover | E value | Per. Ident | Accession |
|---|---|---|---|---|---|---|
| Akkermansia muciniphila strain EB-AMDK-3 chromosome, complete genome | 7755 | 23258 | 98% | 0.0 | 99.53% | CP024738.1 |
| Akkermansia muciniphila strain EB-AMDK-4 chromosome, complete genome | 7751 | 23254 | 98% | 0.0 | 99.51% | CP024740.1 |
| Akkermansia muciniphila strain CBA5201 chromosome, complete genome | 7751 | 23254 | 98% | 0.0 | 99.51% | CP033388.1 |
| Akkermansia muciniphila strain EB-AMDK-7 chromosome, complete genome | 7751 | 23248 | 98% | 0.0 | 99.51% | CP025823.1 |
| Akkermansia muciniphila ATCC BAA-835, complete genome | 7751 | 23254 | 98% | 0.0 | 99.51% | CP001071.1 |
| Akkermansia muciniphila strain EB-AMDK-16 chromosome, complete genome | 7749 | 23243 | 98% | 0.0 | 99.51% | CP025831.1 |
| Akkermansia muciniphila strain H2 chromosome | 7745 | 22576 | 98% | 0.0 | 99.48% | CP010553.1 |

### ➢*Faecalibacterium prausnitzii* - rrn; length = 3,986 nt; coverage = 1000 reads (high)

| Description | Max Score | Total Score | Query Cover | E value | Per. Ident | Accession |
|---|---|---|---|---|---|---|
| Faecalibacterium prausnitzii strain Indica chromosome, complete genome | 6916 | 45825 | 99% | 0.0 | 98.21% | CP023819.1 |
| Faecalibacterium prausnitzii strain A2165 chromosome, complete genome | 6902 | 41296 | 99% | 0.0 | 98.14% | CP022479.1 |
| Faecalibacterium prausnitzii strain APC918/95b chromosome, complete genome | 6859 | 41078 | 99% | 0.0 | 97.96% | CP030777.1 |
| Faecalibacterium prausnitzii strain 942/30-2 chromosome, complete genome | 6735 | 40347 | 99% | 0.0 | 97.39% | CP026548.1 |
