## Supplementary figures and images for "Species- and strain-level assessment using *rrn* long-amplicons suggests donor’s influence on gut microbial transference via fecal transplants in metabolic syndrome subjects"

### Figure S2

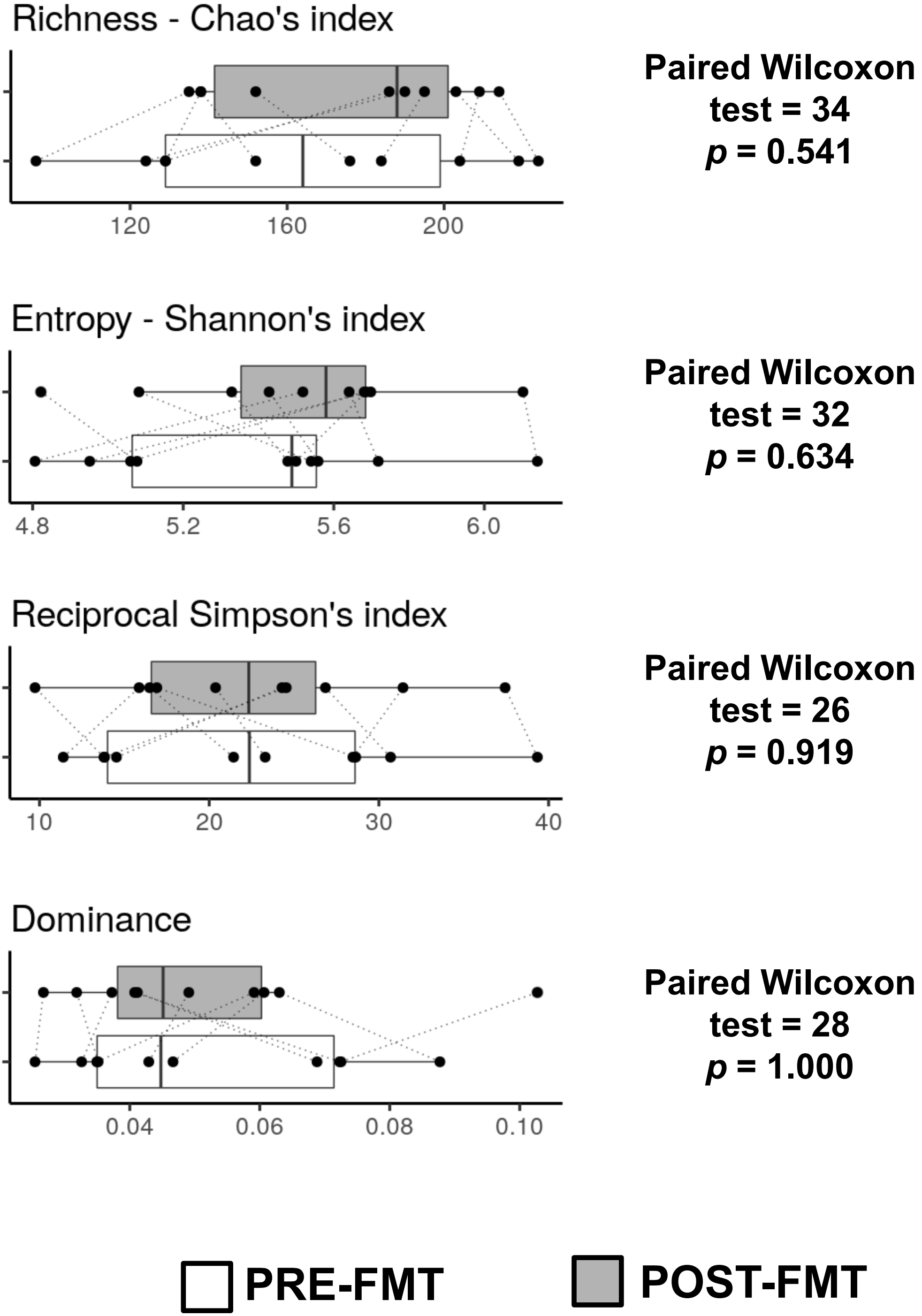
