## Supplementary material for "Species- and strain-level assessment using *rrn* long-amplicons suggests donor’s influence on gut microbial transference via fecal transplants in metabolic syndrome subjects": Figure S3

### *Parabacteroides merdae* SNV-3527 as query (pos 3470-3570)

**Sequences producing significant alignments** Download New Select columns Show 100

select all 5 sequences selected GenBank Graphics Distance tree of results New MSA Viewer

| | Description | Scientific Name | Max Score | Total Score | Query Cover | E value | Per. Ident | Acc. Len | Accession |
|---|---|---|---|---|---|---|---|---|---|
| ☑ | Parabacteroides merdae strain CL06T03C08 chromosome, complete genome | Parabacteroides merdae | 187 | 1125 | 100% | 1e-43 | 100.00% | 4697288 | CP072229.1 |
| ☑ | Uncultured bacterium clone LM0ACA6ZA05RM1 genomic sequence | uncultured bacterium | 182 | 182 | 100% | 5e-42 | 99.01% | 706 | EU066707.1 |
| ☑ | Uncultured bacterium clone LM0ACA6ZD02FM1 genomic sequence | uncultured bacterium | 182 | 182 | 100% | 5e-42 | 99.01% | 710 | EU063890.1 |
| ☑ | Uncultured bacterium clone LM0ACA4ZC10FM1 genomic sequence | uncultured bacterium | 182 | 182 | 100% | 5e-42 | 99.01% | 585 | EU063717.1 |
| ☑ | Bacteroides merdae 23S ribosomal RNA gene, partial sequence | Parabacteroides merdae ATCC 43184 | 182 | 182 | 100% | 5e-42 | 99.01% | 574 | AY155595.1 |

Download GenBank Graphics Sort by: E value

Parabacteroides merdae strain CL06T03C08 chromosome, complete genome

Sequence ID: CP072229.1 Length: 4697288 Number of Matches: 6

Range 1: 37534 to 37634 GenBank Graphics ▼ Next Match ▲ Previous Match

| Score | Expect | Identities | Gaps | Strand |
|---|---|---|---|---|
| 187 bits(101) | 1e-43 | 101/101(100%) | 0/101(0%) | Plus/Minus |

```
Query  1      CCTGATTACGTCCATTTTGCCTTGTGCAAGCACGCGGCATACTATCAAGGTCGACTCTCC  60
              ||||||||||||||||||||||||||||||||||||||||||||||||||||||||||||
Sbjct  37634  CCTGATTACGTCCATTTTGCCTTGTGCAAGCACGCGGCATACTATCAAGGTCGACTCTCC  37575

Query  61     CTGCGGATTTGCCTACAGGAATCTACATCTACACTCTTCAA  101
              |||||||||||||||||||||||||||||||||||||||||
Sbjct  37574  CTGCGGATTTGCCTACAGGAATCTACATCTACACTCTTCAA  37534
```

Strain CL06T03C08 genotype A3527

Download GenBank Graphics

Uncultured bacterium clone LM0ACA6ZA05RM1 genomic sequence

Sequence ID: EU066707.1 Length: 706 Number of Matches: 1

Range 1: 210 to 310 GenBank Graphics ▼ Next Match ▲ Previous Match

| Score | Expect | Identities | Gaps | Strand |
|---|---|---|---|---|
| 182 bits(98) | 5e-42 | 100/101(99%) | 0/101(0%) | Plus/Minus |

```
Query  1    CCTGATTACGTCCATTTTGCCTTGTGCAAGCACGCGGCATACTATCAAGGTCGACTCTCC  60
            ||||||||||||||||||||||||||||||||||||||||||| ||||||||||||||||
Sbjct  310  CCTGATTACGTCCATTTTGCCTTGTGCAAGCACGCGGCATACTGTCAAGGTCGACTCTCC  251

Query  61   CTGCGGATTTGCCTACAGGAATCTACATCTACACTCTTCAA  101
            |||||||||||||||||||||||||||||||||||||||||
Sbjct  250  CTGCGGATTTGCCTACAGGAATCTACATCTACACTCTTCAA  210
```

Strain LM0ACA6ZA05RM1 genotype G3527

Download GenBank Graphics

Uncultured bacterium clone LM0ACA6ZD02FM1 genomic sequence

Sequence ID: EU063890.1 Length: 710 Number of Matches: 1

Range 1: 316 to 416 GenBank Graphics ▼ Next Match ▲ Previous Match

| Score | Expect | Identities | Gaps | Strand |
|---|---|---|---|---|
| 182 bits(98) | 5e-42 | 100/101(99%) | 0/101(0%) | Plus/Plus |

```
Query  1    CCTGATTACGTCCATTTTGCCTTGTGCAAGCACGCGGCATACTATCAAGGTCGACTCTCC  60
            ||||||||||||||||||||||||||||||||||||||||||| ||||||||||||||||
Sbjct  316  CCTGATTACGTCCATTTTGCCTTGTGCAAGCACGCGGCATACTGTCAAGGTCGACTCTCC  375

Query  61   CTGCGGATTTGCCTACAGGAATCTACATCTACACTCTTCAA  101
            |||||||||||||||||||||||||||||||||||||||||
Sbjct  376  CTGCGGATTTGCCTACAGGAATCTACATCTACACTCTTCAA  416
```

Strain LM0ACA6ZD02FM1 genotype G3527

Download GenBank Graphics

Uncultured bacterium clone LM0ACA4ZC10FM1 genomic sequence

Sequence ID: EU063717.1 Length: 585 Number of Matches: 1

Range 1: 396 to 496 GenBank Graphics ▼ Next Match ▲ Previous Match

| Score | Expect | Identities | Gaps | Strand |
|---|---|---|---|---|
| 182 bits(98) | 5e-42 | 100/101(99%) | 0/101(0%) | Plus/Plus |

```
Query  1    CCTGATTACGTCCATTTTGCCTTGTGCAAGCACGCGGCATACTATCAAGGTCGACTCTCC  60
            ||||||||||||||||||||||||||||||||||||||||||| ||||||||||||||||
Sbjct  396  CCTGATTACGTCCATTTTGCCTTGTGCAAGCACGCGGCATACTGTCAAGGTCGACTCTCC  455

Query  61   CTGCGGATTTGCCTACAGGAATCTACATCTACACTCTTCAA  101
            |||||||||||||||||||||||||||||||||||||||||
Sbjct  456  CTGCGGATTTGCCTACAGGAATCTACATCTACACTCTTCAA  496
```

Strain LM0ACA4ZC10FM1 genotype G3527

Download GenBank Graphics

Bacteroides merdae 23S ribosomal RNA gene, partial sequence

Sequence ID: AY155595.1 Length: 574 Number of Matches: 1

Range 1: 391 to 491 GenBank Graphics ▼ Next Match ▲ Previous Match

| Score | Expect | Identities | Gaps | Strand |
|---|---|---|---|---|
| 182 bits(98) | 5e-42 | 100/101(99%) | 0/101(0%) | Plus/Minus |

```
Query  1    CCTGATTACGTCCATTTTGCCTTGTGCAAGCACGCGGCATACTATCAAGGTCGACTCTCC  60
            ||||||||||||||||||||||||||||||||||||||||||| ||||||||||||||||
Sbjct  491  CCTGATTACGTCCATTTTGCCTTGTGCAAGCACGCGGCATACTGTCAAGGTCGACTCTCC  432

Query  61   CTGCGGATTTGCCTACAGGAATCTACATCTACACTCTTCAA  101
            |||||||||||||||||||||||||||||||||||||||||
Sbjct  431  CTGCGGATTTGCCTACAGGAATCTACATCTACACTCTTCAA  391
```

Strain ATCC43184 genotype G3527
