## Supplementary material for "Species- and strain-level assessment using *rrn* long-amplicons suggests donor’s influence on gut microbial transference via fecal transplants in metabolic syndrome subjects": Figure S4

Albacore v2.1.3 | Guppy v5.0.11

### Run1 – Per base sequence quality

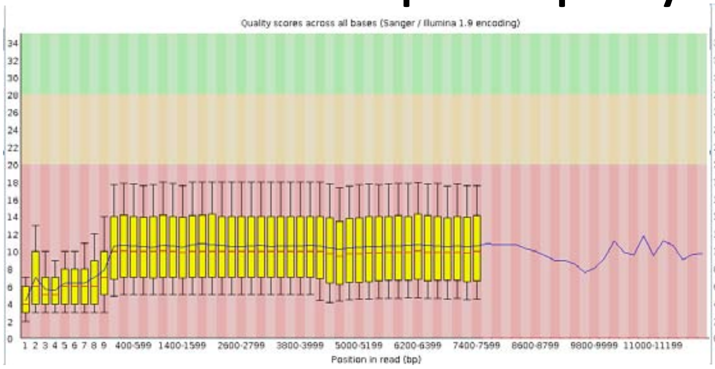


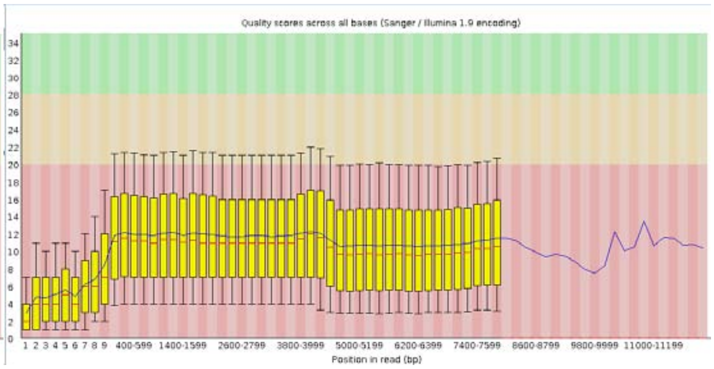


### Run1 – Per sequence quality scores

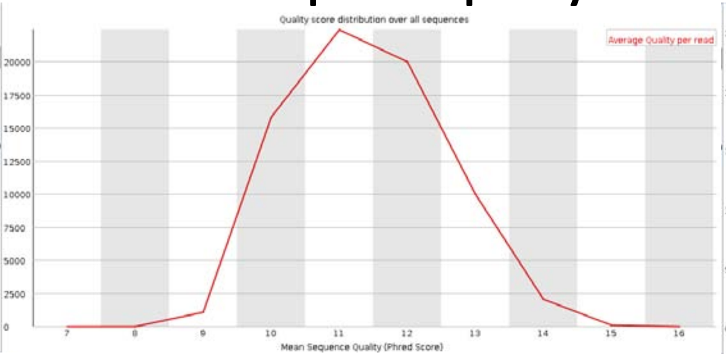


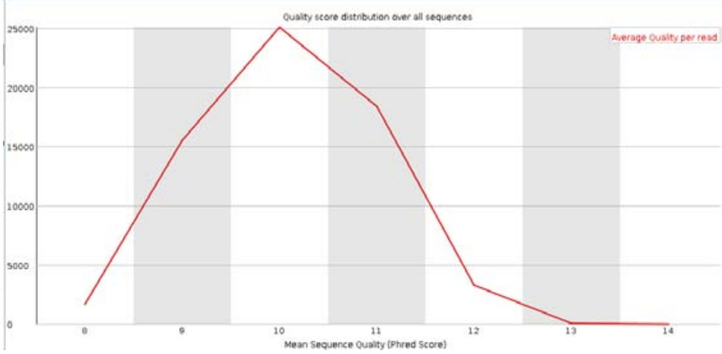


### Run1 – Per base sequence content

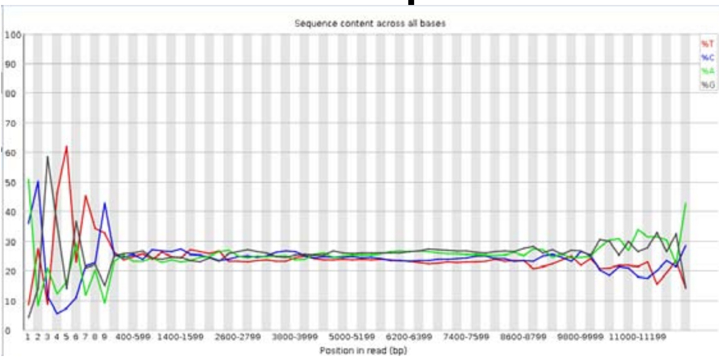


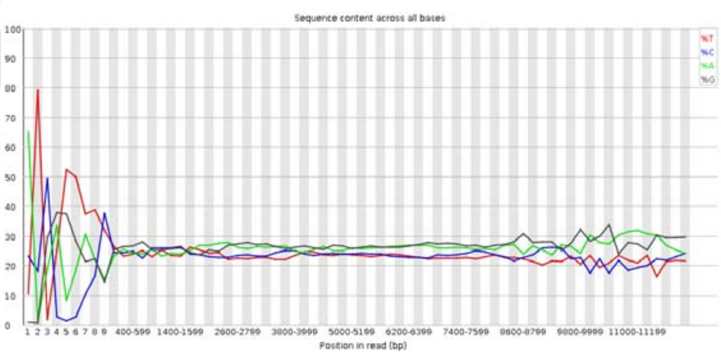


### Run1 – Sequence length distribution

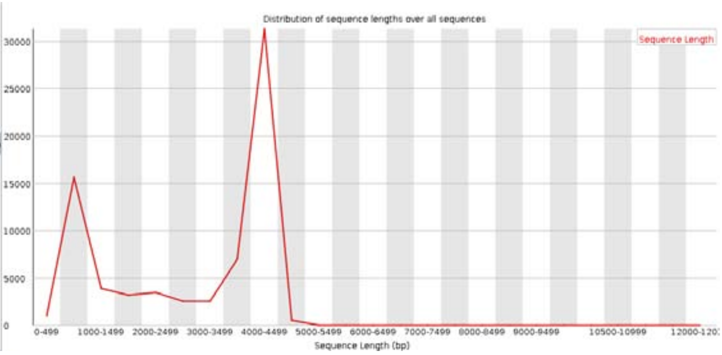


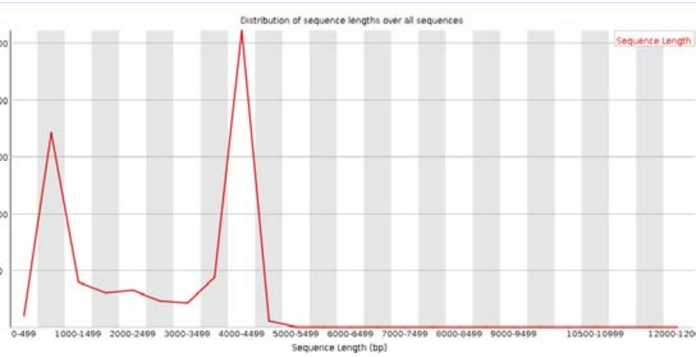

Albacore v2.1.3 Guppy v5.0.11

### Run2 – Per base sequence quality

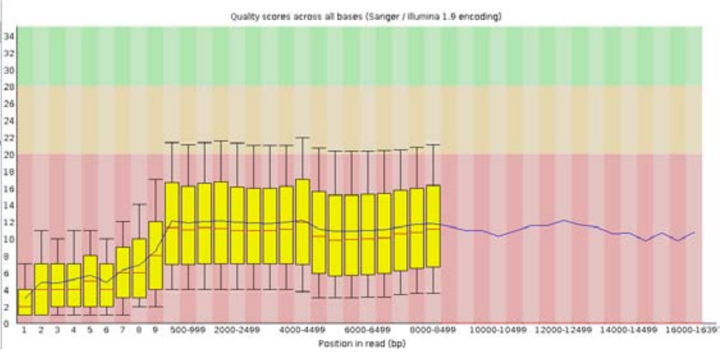


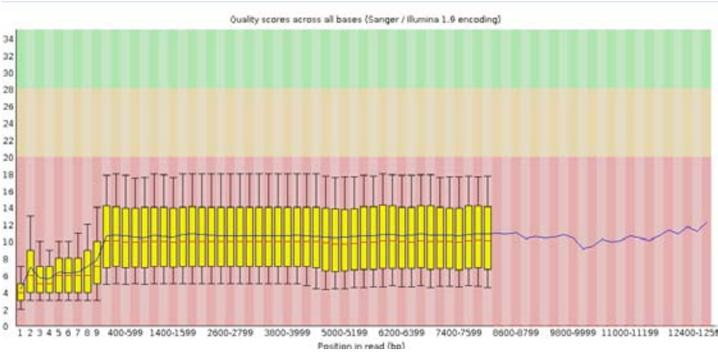


### Run2 – Per sequence quality scores

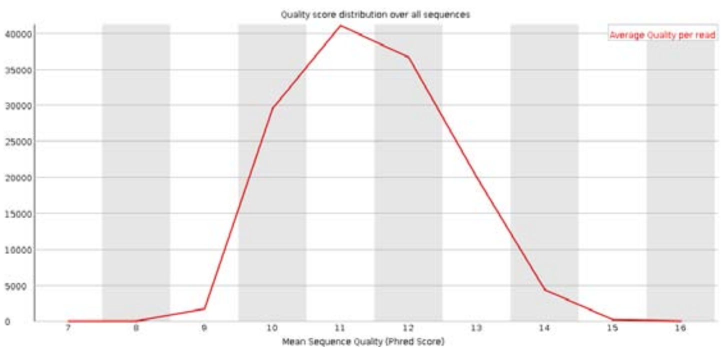


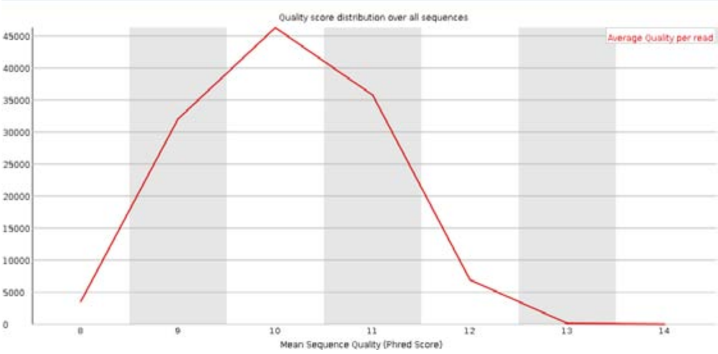


### Run2 – Per base sequence content

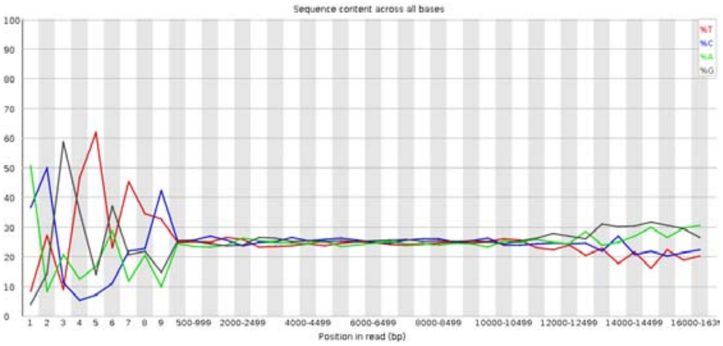


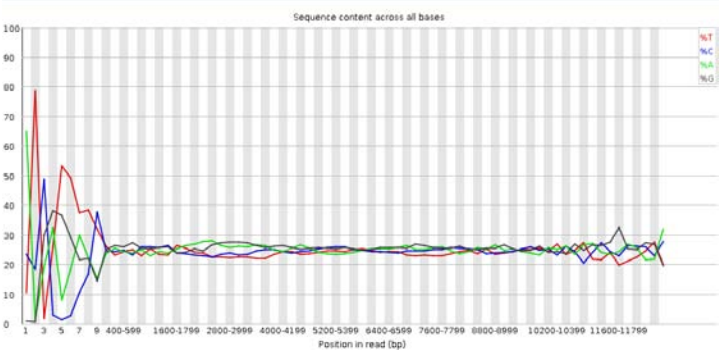


### Run2 – Sequence length distribution

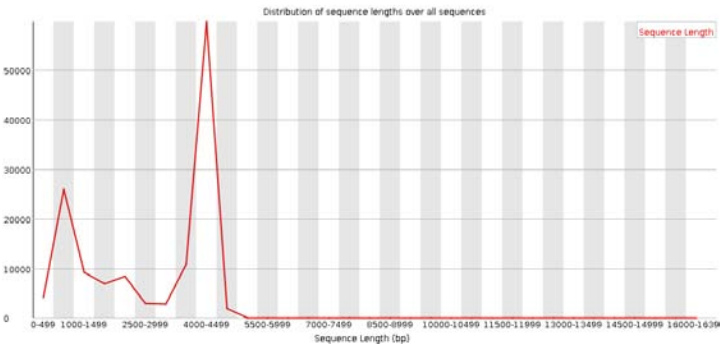


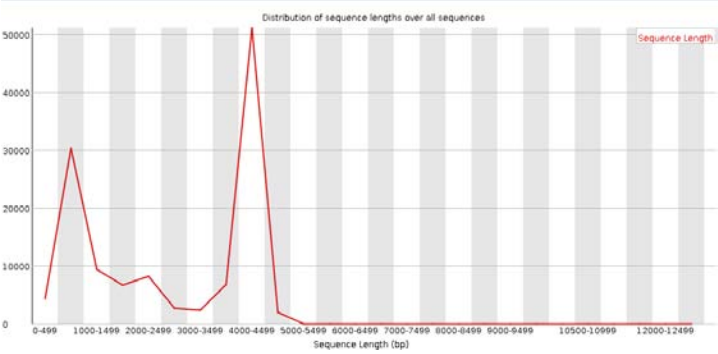

Albacore v2.1.3 Guppy v5.0.11

### Run3 – Per base sequence quality

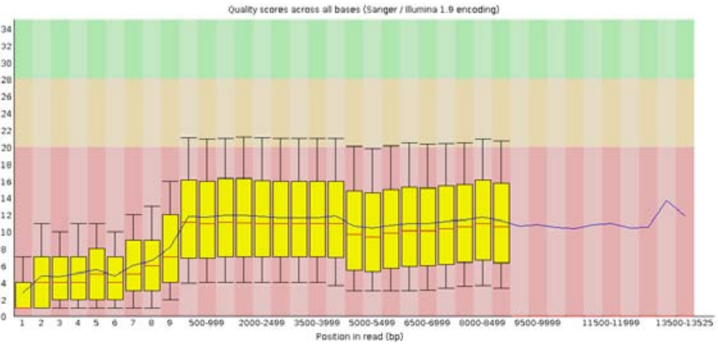


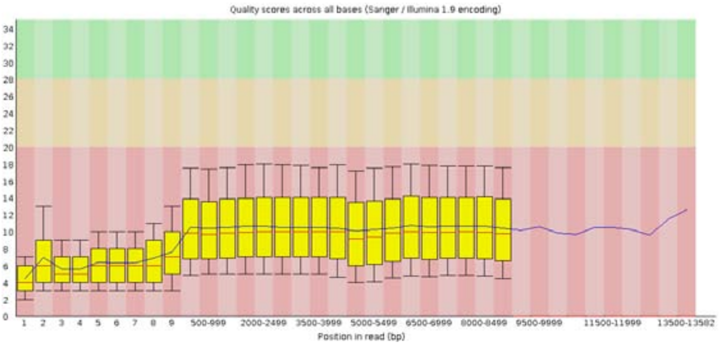


### Run3 – Per sequence quality scores

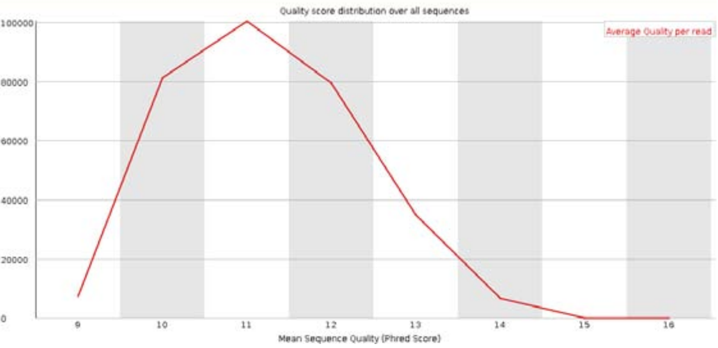


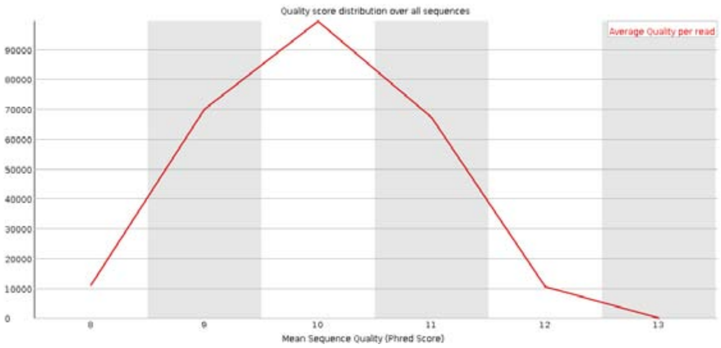


### Run3 – Per base sequence content

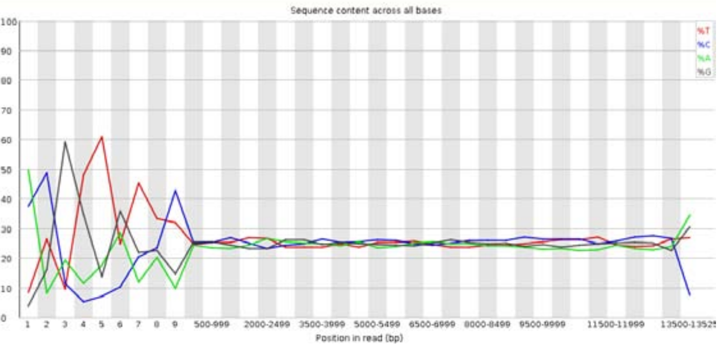


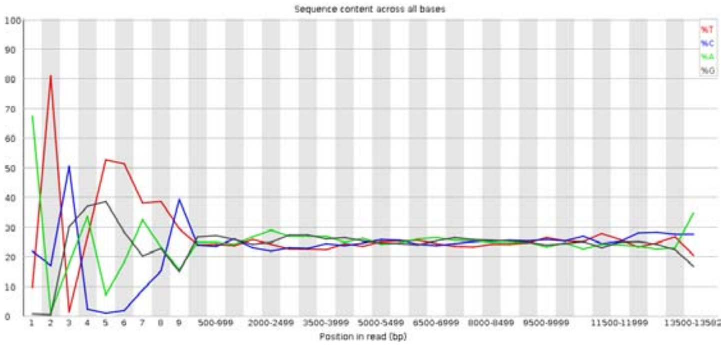


### Run3 – Sequence length distribution

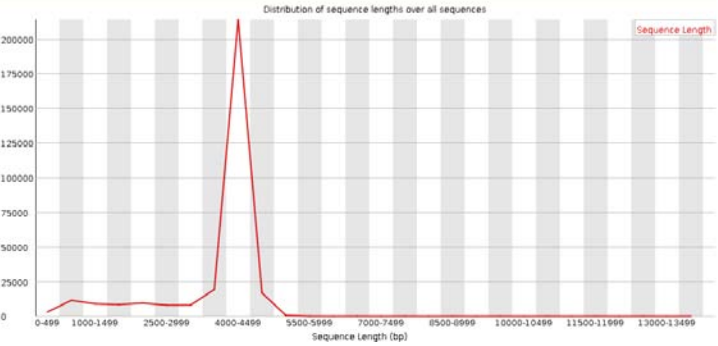


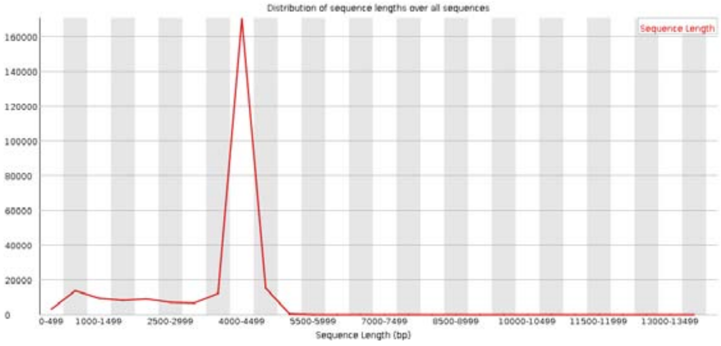
